# Effects of Endophytic Fungus MF23 on *Dendrobium nobile* Lindl. in an artificial primary environment

**DOI:** 10.1101/2019.12.16.878066

**Authors:** Xiaolin Xu, Qing Li, Gang Ding, Biao Li, Shunxing Guo

## Abstract

The quality of *Dendrobium nobile* Lindl. is related to the endophytic fungus. It had reported that the mycorrhizal fungus MF23 helps to increase the content of dendrobine, but few studies had explained the mechanism of the phenomenon. In previous study, we verified the symbiotic mechanism of the mycorrhizal fungus MF23 with *D. nobile* on the agar medium. In this study, the research carried on the bark medium, nearly like the natural environment, which had special meaning because of its benefits for the widely application. As a result, it showed a significant effect especially in the later period of the cultivation, suggesting that mycorrhizal fungus MF23 had a promotion for *D. nobile* in the natural environment, which enabled the application of the technique in the experimental field. It also implied that post-modification enzyme genes might play an important role in stimulating biosynthesis of dendrobine.

## 1. Introduction

*Dendrobium nobile* Lindl. is a traditional medicinal plant which has the reputation of “fairy grass”. With the effect of maintaining gastric tonicity, enhancing production of body fluid, relieving symptom of dryness and curing the symptom of body heat [1], it has been widely used in the clinical and Chinese medicine prescription. Dendrobine, a sesquiterpene compound, is considered as the effective component of *D. nobile*, the Pharmacopoeia of the People’s Republic of China regards it as an important indicator of the quality of *D. nobile.* The main active ingredients of *D. nobile* include dendrobine, polysaccharides, flavonoids, bibenzyls and so on [2–4]. It had reported that *D. nobile* has some pharmacological activities such as antioxidant, antithrombotic and anti-Alzheimer’s disease [5–6].

The growth of the Orchidaceae is closely related to the effect of mycorrhizal fungus [7], it had reported that the fungi could promote the germination of the orchid species [8]. As we all know, Orchidaceae seeds, only with undifferentiated embryos, are small as dust. In this case, they must rely on fungal infection to provide nutrients to germinate under natural conditions. Guo *et al* [9–10] obtained a number of fungi from the Dendrobium, which could effectively promote the germination of Dendrobium seeds. Fungi also affected the growth of the plant [11], through the way of invading the root cells, and forming mycorrhizal symbioses with the root. It could supply the water and nutrients such as nitrogen and phosphorus to the host plant in a further thought [12]. Up to date, few studies have been published referring to the stress resistance of medicinal plant., one of which had shown that it could promote the stress resistance with the effect of mycorrhizal fungus. The fungus elicitor of the genus could induce the activity of lipoxygenase, peroxidase and phenylalanine ammonia lyase in the protocorm of *D. candidum* to protect itself, which were involved in the synthesis of resistant substances [13]. The effect of endophytic fungus on host plant has been examined in more details. It showed effective increase of the active ingredients content such as steroids and alkaloids after the infection [14–15].

MF23, belongs to *Mycena* sp., is a mycorrhizal fungus separated from *D. officinale* Kimura et Migo. It had reported that MF23 had the function of promoting the growth of Dendrobium plant [16–17]. In recent studies, it proved that MF23 could promote the growth of the *D. officinale*, and increase the dendrobine content of *D. nobile* [18–19]. However, these researches carried out on the base of agar medium with rich nutrition. Our previous research [20] had been carried out with the agar medium, with good repeatability and controllability for the analysis of the results. However, it was not matched with the natural environment of Dendrobium, surroundings with stones or barks lacking of necessary nutrition. In this case, in order to make the results of research apply for the production and application in reality, this study had performed on the chestnut bark, offering a general natural environment to study the effect of MF23 on *D. nobile*, to provide basic materials and evidence for the MF23 induced symbiotic growth promoting mechanism in natural environment. The purpose of the study is to explore the effect and the mechanism of mycorrhizal fungi on the *D. nobile* with bark medium, and lay a foundation for the possibility and mechanism of using mycorrhizal fungi to promote the production of *D. nobile* in the field.

## 2. Results

### 2.1. The effect of MF23 on the growth state of D. nobile

The survival rate of *D. nobile* has a decreased trend on the total level both in the experimental and experimental groups, but with the extension of the time after cultivation, the experimental groups had a higher survival rate than the control groups, and maintain a relatively stable trend in the period of inoculation. At 12^th^ week, the survival rate had a significant difference between the control and the experimental group, and at 24^th^ week, the survival rate of experimental group was only 72.9% whereas the experimental group had higher rate, more than 85.4%, which formed a striking contrast with the control group (**Figure 1A**). In addition, the stem diameter of *D. nobile* presented a significant difference between the experimental and the control group since the 9^th^ week (**Figure 1B**), In view of the weight of the fresh stem, the experimental groups showed significant promotion than the control groups since the 3^rd^ week (**Figure 1C**). In general, the MF23 did promote the growth of the *D. nobile* according to the results.

**Figure 1.**
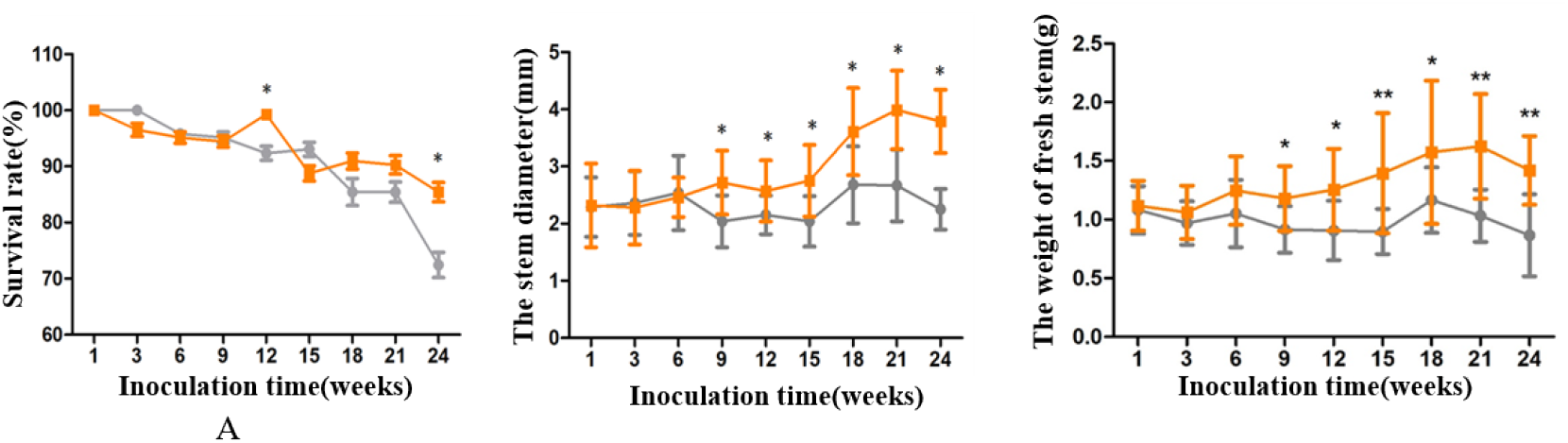
The changes of morphology on *D. nobile* grown on the bark medium between the control (grey line) and the experimental group (color line). Values are presented as the means ±SD (n=12); values with * are of statistical significance at P < 0.05 A: the comparison of survival rate B: the comparison of stem diameter C: the comparison of weight of fresh stem

### 2.2. The effect of MF23 on the content of dendrobine

In general, the dendrobine content had a significant difference between the experimental and the control groups. It was not so significant in the early six weeks, but as time goes on, it showed that the dendrobine had been accumulated obviously both in the control and the experimental group (**Figure 2**), especially at the 21^st^ week, the obvious difference of the debdrobine content occurred in the experimental group (P<0.01), reaching to 0.146%, which is almost three times more than that in the control groups (0.051%).

**Figure 2.**
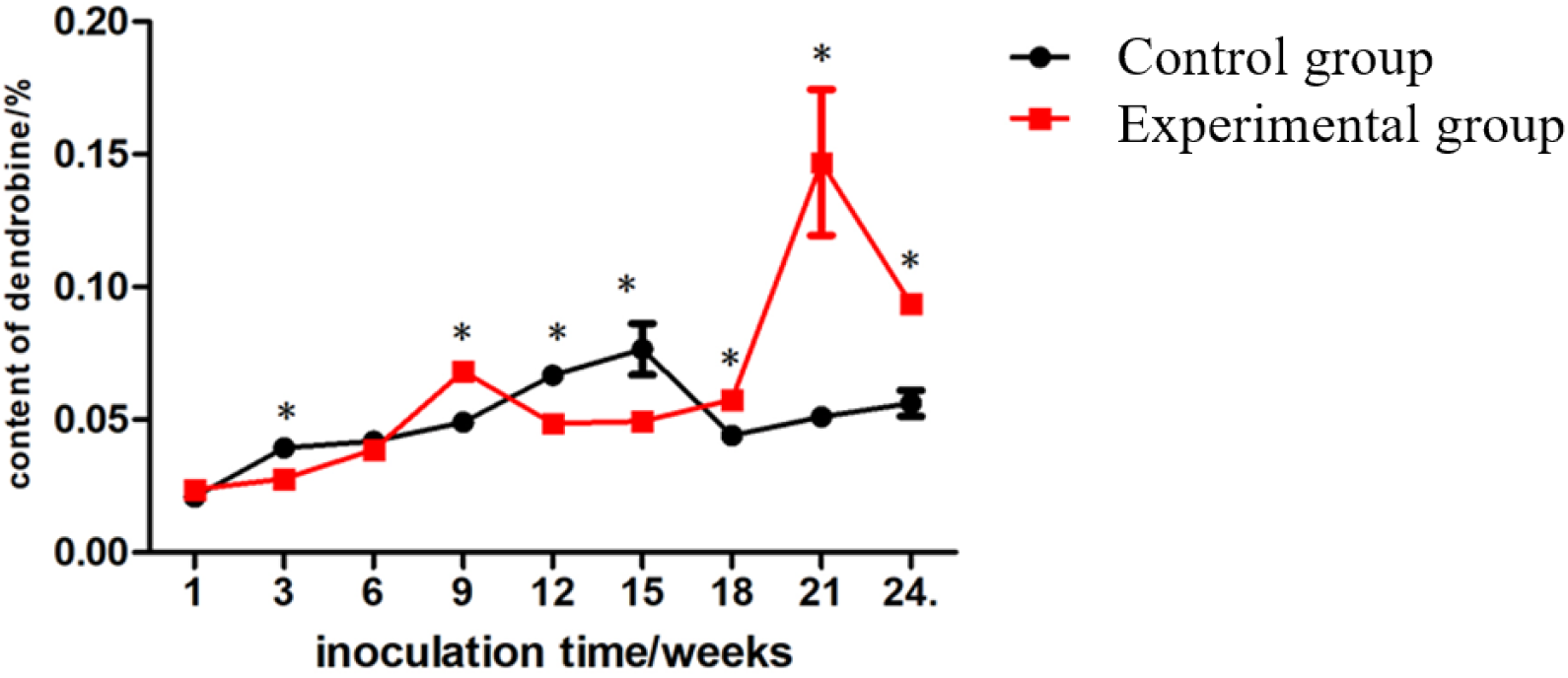
The difference of the dendrobine content between the control and the experimental group. Values are presented as the means ±SD (n=3); values with * are of statistical significance at P < 0.05

### 2.3. The infection process of MF23 on roots of D. nobile

Hyphae were present in the velamen at 6^th^ week after inoculation (**Figure 3A**), then occurred in the exodermis by 9 weeks (**Figure 3B**), while appeared in the cortex by 15 weeks (**Figure 3C**) and in passage cells by 21 weeks (**Figure 3D**). Also, it showed that the mycelium in the cortex of *D. nobile* was densely covered at 18^th^ week, but rarely seen by the 21^st^ and 24^th^ week.

**Figure 3.**
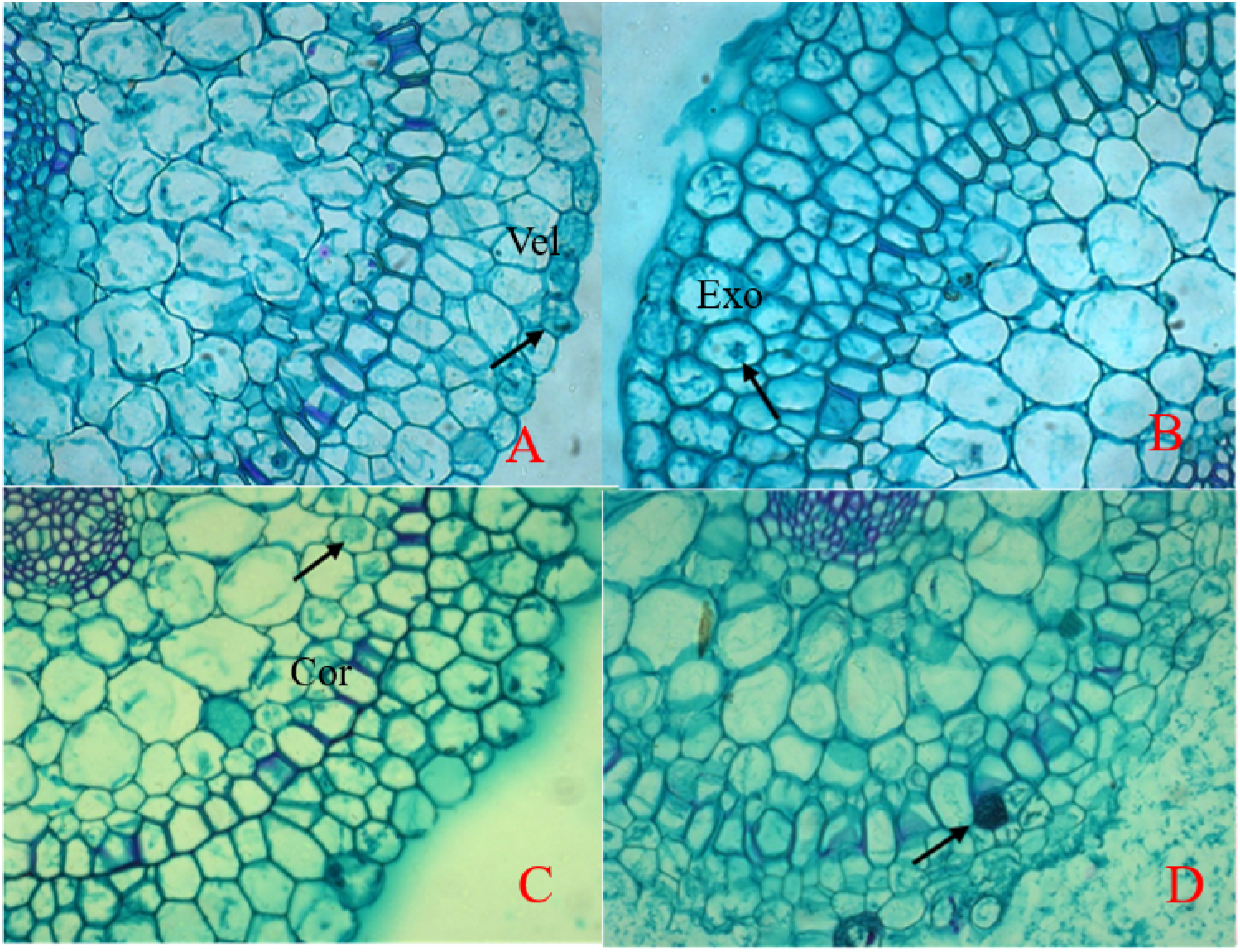
Light micrographs of interactions between the roots of *D. nobile* and MF23 (A-D) A: Mycelium entered into velamen of the root after 6 weeks (×20). B: Mycelium entered into exodermis of the root after 9 weeks (×20). C: Mycelium entered into cortex of the root after 15 weeks (×20). D: The hyphal coils appeared after 21 weeks (×20). (vel represents velamen; exo represents exodermis; cor represents cortex.)

**Figure 4.**
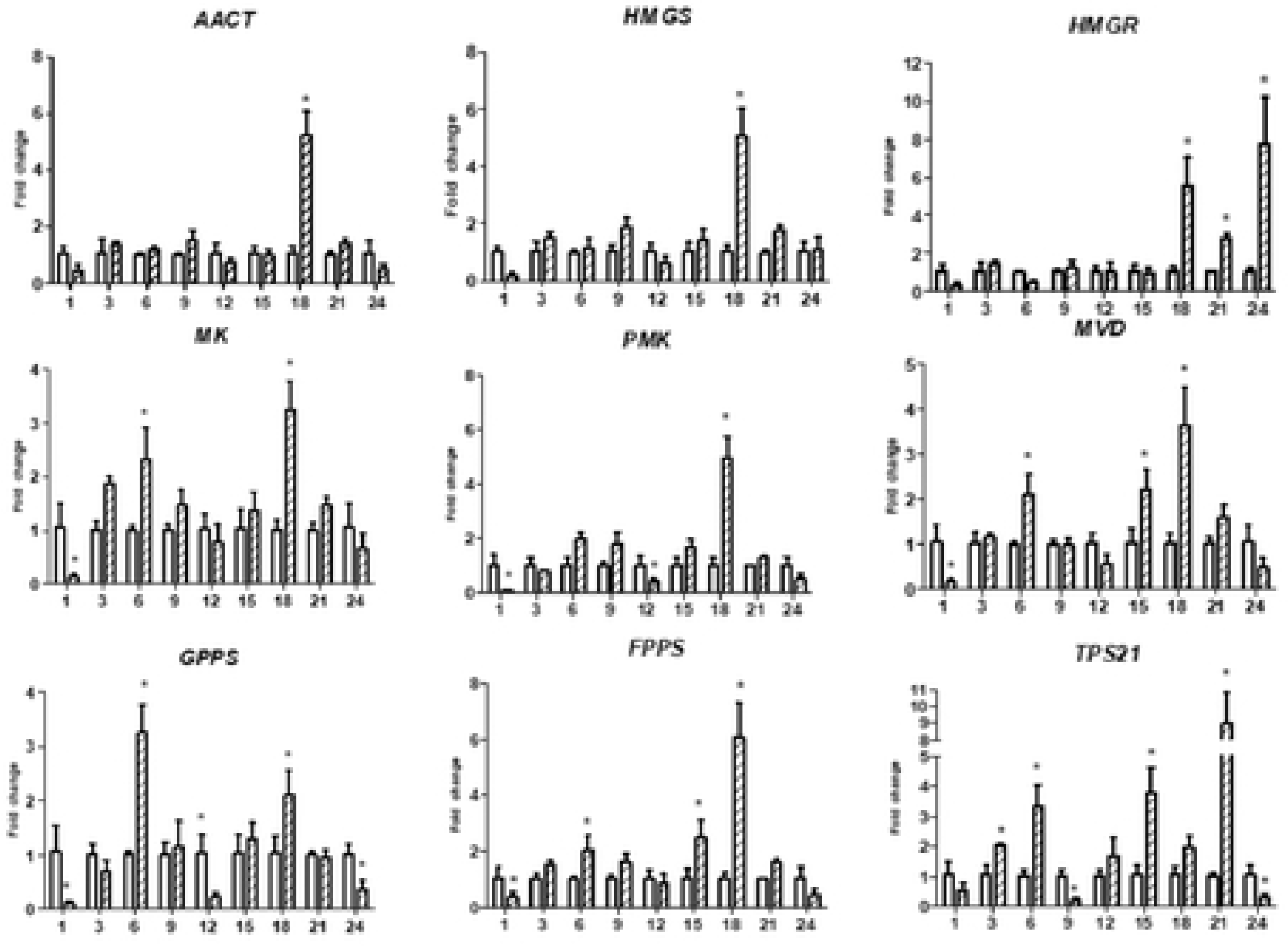
qRT-PCR analysis of key enzyme-coding genes involved in the MVA pathway in *D. nobile*. The y-axes correspond to the mean fold changes in expression values, and the x-axes display the times of Symbiotic culture (/weeks). White bars represent the control group, and shadow bars represent the experimental group. For each qRT-PCR validation, three technical replicates were used, with a minimum of three biological replicates.

**Figure 5.**
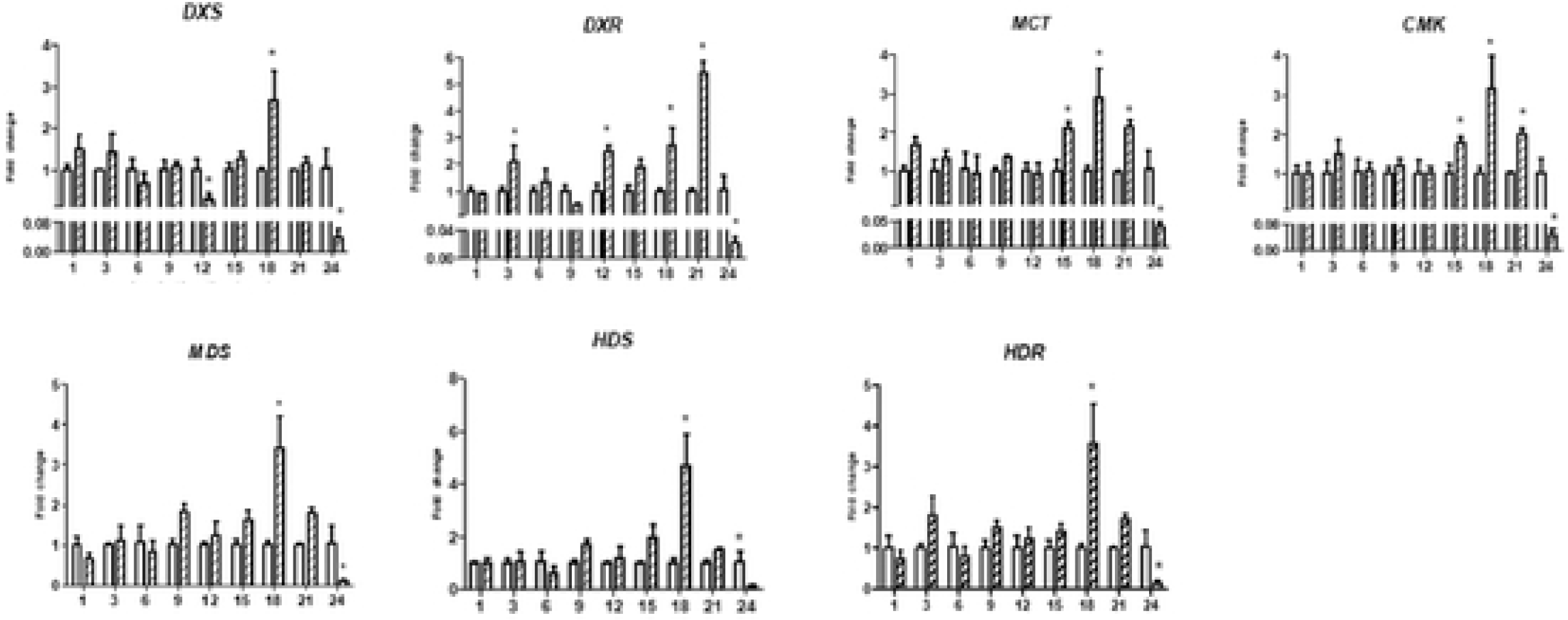
qRT-PCR analysis of key enzyme-coding genes involved in the MEP pathway in *D. nobile*. The y-axes correspond to the mean fold changes in expression values, and the x-axes display the times of symbiotic culture (/weeks). White bars represent the control group, and shadow bars represent the experimental group. For each qRT-PCR validation, three technical replicates were used, with a minimum of three biological replicates.

**Figure 6.**
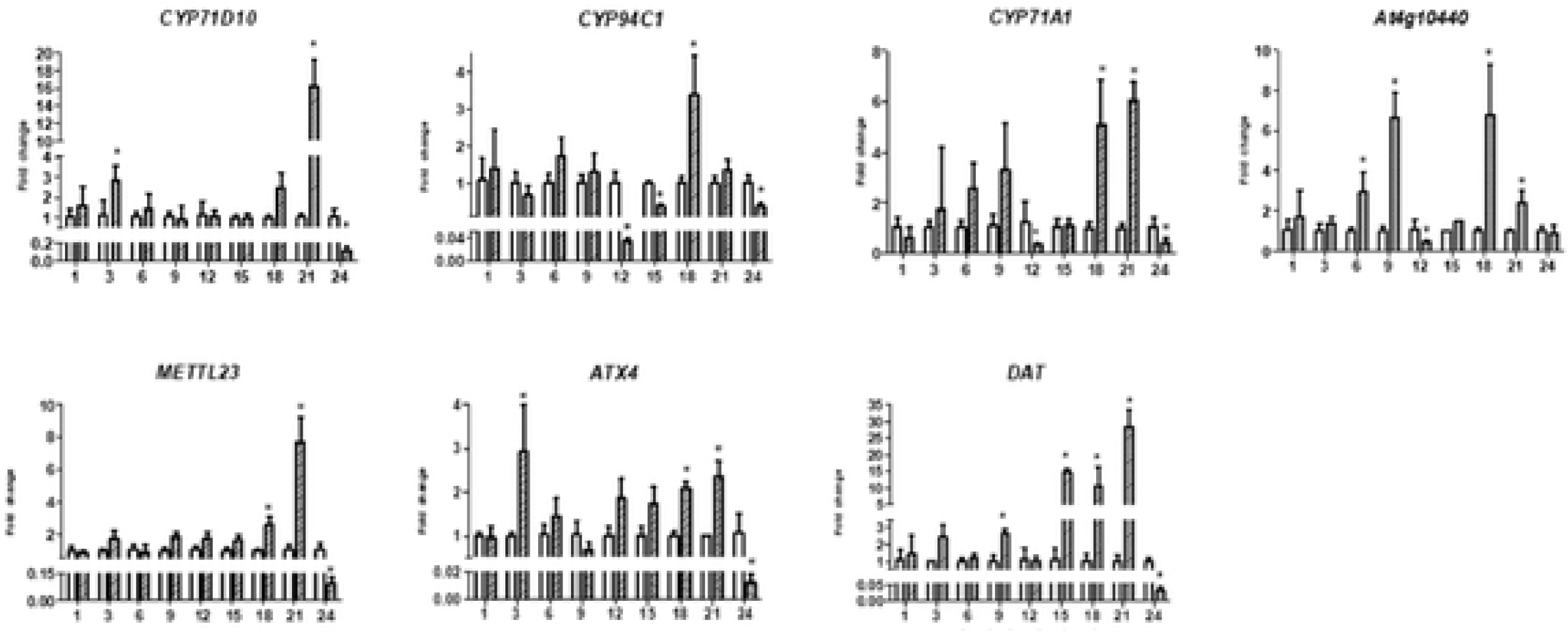
qRT-PCR analysis of key enzyme-coding genes involved in post-modification in *D. nobile*. The y-axes correspond to the mean fold changes in expression values, and the x-axes display the times of symbiotic culture (/weeks). White bars represent the control group and shadow bars represent the experimental group. For each qRT-PCR validation, three technical replicates were used, with a minmum of three biological replicates.

**Figure 7.**
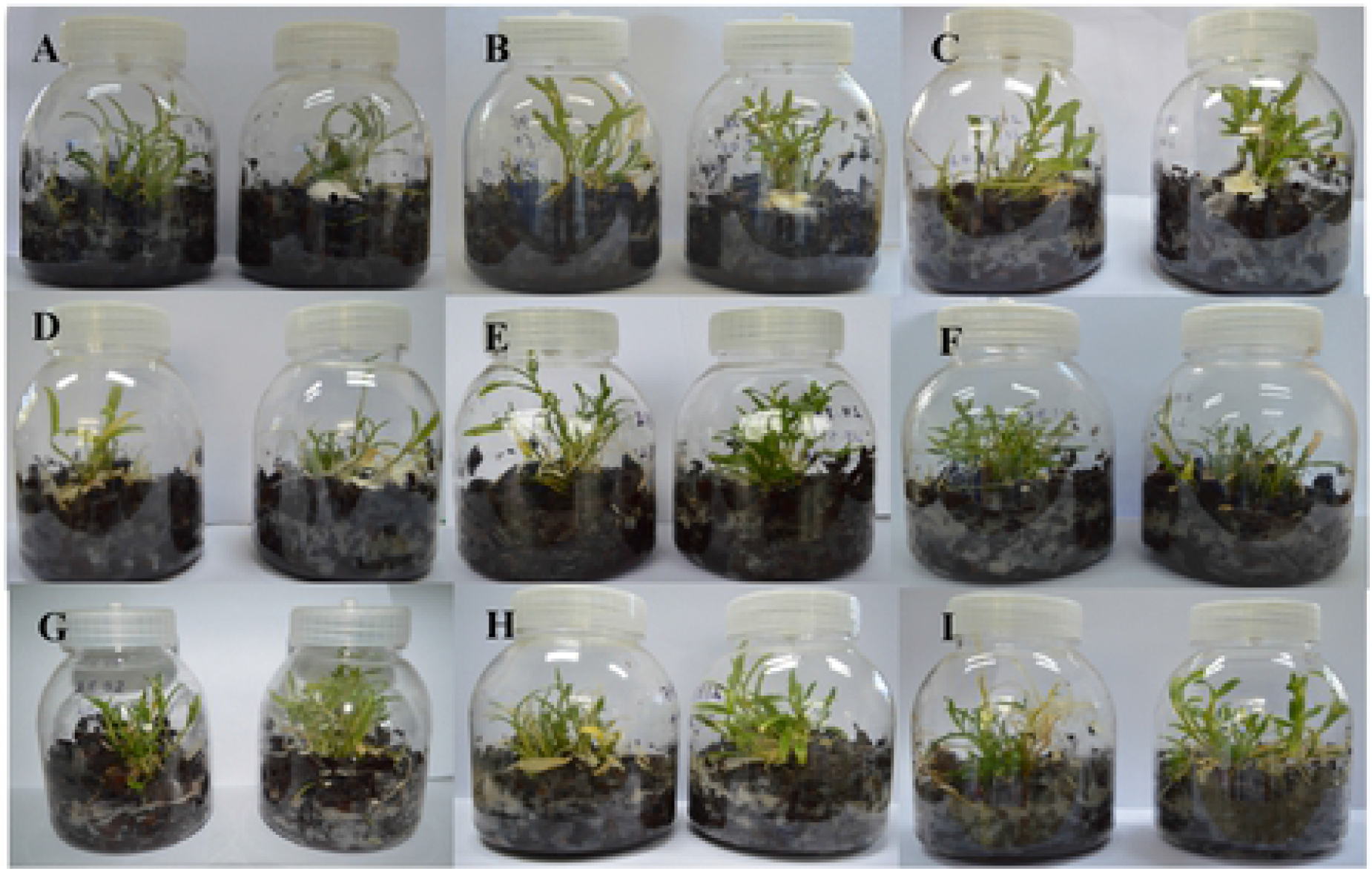
The effect of MF23 on the growth morphology of *D. nobile* cultivated inbark medium. (A-I represent cultivation after 1, 3, 6, 9, 12, 15, 18, 21, 24 weeks, the left ones are control groups, the right ones are expression groups respectively)

### 2.4. Validation and expression analysis of genes involved in the formation of the backbone of the sesquiterpene alkaloid dendrobine

The key enzyme genes involved in the upstream synthetic pathway of dendrobine were picked out for a qRT-PCR analysis. It could be seen from the figure (**Figure S1**) that genes *MK*, *MVD*, *GPPS*, *FPPS* and *TPS21*were up-regulated at early weeks (6 weeks), *MVD*, *FPPS* and *TPS21*were up-regulated at middle weeks (15 weeks), and all the genes were up-regulated at 18^th^ week, but dropped to a lower expression level and had no significant difference compared with the control group at 21^st^ week, the time when the experimental group reached the maximum content of dendrobine. At the 24^th^ week, the expression levels of all the key enzyme genes in the mevalonate (MVA) pathway were down-regulated except *HMGR*.

The expression pattern of key enzyme genes in the methyl-D-erythritol 4-phosphate (MEP) pathway (**Figure S2**) was almost the same as that in the control group in prior inoculation stage, only the gene DXR was up-regulated at early period (6 weeks). At the middle period (18 weeks), all the genes (*DXS*, *DXR*, *MCT*, *CMK*, *MDS*, *HDS* and *HDR*) were up-regulated, and the activation was much greater than any other times. However, during the late stage of inoculation (24^th^ week), the expression pattern of key enzyme genes was similar to that of MVA pathway, down-regulated to a low level.

### 2.5. Validation and expression analysis of genes involved in post-modification

The expression of genes involved in post-modification process were more consistent with the change of the dendrobine content (**Figure S3**). At early weeks (6 weeks), genes *CYP71D10*, *At4g10440* and *ATX4* were up-regulated, and genes had no significant change during the middle period. All the genes exhibited higher expression levels at 18 weeks after inoculation. Their levels of expression were higher than the control groups, reaching to the significant difference at 21 weeks, and almost all the genes except *CYP94C1*, *At4g10440 and ATX4* exhibited higher expression levels after inoculation than any other stages, which was consistent with the maximizing of dendrobine content. In conclusion, the enzyme genes involved in post-modification had closer relationship with the content of dendrobine.

## 3. Discussion

### 3.1. MF23 had significant effect on the growth state of the D. nobile on the bark medium

It showed that MF23 had significant effect on the growth of the *D. nobile* on the bark medium. It could be found that the time point at which the growth index of *D. nobile* in the experimental group began to increase significantly (9^th^ week) was highly consistent with the time when the MF23 hyphae began to enter the interior of the root. It can be inferred that the increase in growth of *D. nobile* is closely related to the infection of hyphae. As early as 1983, Ames *et al* [21] used radioisotope labeling technology to find that AM fungal hyphae can absorb nitrogen from soil outside the root and transport it to the root of the host. In addition, Li *et al* [22] had proved that compared with the root system of plants, the absorption area of extra-root hyphal networks is wider, which is more conducive to the absorption of water and a large number of mineral elements. Moreover, endophytic fungi can increase the pH of the mycelium by secreting protons and organic acids to activate substances such as slightly soluble phosphates in the matrix, so that nutrients such as phosphate can be increased in the rhizosphere [23]. This is probably one of the important reasons why *D. nobile* can still maintain a good growth situation on nutrient-deficient bark. The dendrobine content began to decrease after 21 weeks inoculation, it is presumed that the fungal mycelium might be consumed by *D. nobile* after 21 weeks, which is involved in the nutrient modes of Orchid plants [24].

### 3.2. Meaningful change trend of dendrobine content on bark mediums

In this study, MF23 showed a significant effect on the content of dendrobine, and the maximum increase was 187.7% compared with the control group. It had reported that there may be a negative feedback regulation mechanism in the synthesis pathway of salicylic acid in *Scutellaria baicalensis* Georgi, which was involved in the accumulation of the secondary metabolite like baicalin and baicalein [25]. It was speculated that salicylic acid may participate in or trigger the regulation process to prevent excessive accumulation of the plant. Some other researches [26–27]also had showed that it mainly existed a feedback inhibition in biosynthesis pathway of plant. Therefore, it could be supposed that after 21 weeks of inoculation, the biosynthetic pathway of the experimental group was negatively regulated and inhibited, leading to the decrease in the content of dendrobine after 24 weeks, which speculated that the secondary metabolites products of the *D. nobile* may keep a balance to protect itself from the risk of too much alkaloids.

### 3.3. Infection progress of MF23 with D. nobile from velamen to exodermis then to cortex

It could be seen that the hyphae of the MF23 were appeared from the velamen to the exodermis then to the cortical cells and gradually advanced, and what is worthy to observe was a small amount of hyphal distribution in non-channel cells. Two reasons might explain this phenomenon, one is that the process of MF23 influx into the cortex from passage cells occurred precisely at the time we did not investigate. Otherwise, passage cells probably not the only channel for MF23 to enter cortical cells. MF23 could pass into the cortex through non-channel cells directly. We could find that the mycelium in the cortex of *D. nobile* densely covered at 18^th^ week, but rarely seen at the 21^st^ and 24^th^ week, and it had reported that Orchidaceae might digest endophytic fungal to obtain nutrients for growth [24]. Therefore, we may regard it as the period for the large-scale digestion of fungal hyphae to achieve energy and nutrients. In addition, it also reported that there were differences in the distribution of hyphae in different parts of the root, Pan *et al* [28] found that most of the Orchidaceae had no mycorrhizal infection at the root tip, in general, it may also be related to the sampling site.

### 3.4. MF23 is effective to the enzyme genes of MVA pathway—the upstream synthetic pathway of dendrobine in natural environment

From the perspective of molecular biology, it showed that the key enzyme gene in the MVA pathway of *D. nobile* grew on the bark medium is effective to the induction of MF23, but it appeared that an unequal relationship existed in the MVA pathway between the biosynthesis of dendrobine and the expression of key enzyme genes. Studies have showed that genes often have an unequal relationship between transcriptome and proteome levels, leading to unequal symmetry with catalytic products, which may be resulted from the enzymatic processing and the change of regulation before and after gene translation [29]. For instance, when *Fusarium* was induced for 2 months, the sesqui-TPS gene was active, but the increase of sesquiterpenoid content was at the next adjacent sampling point, as well as the time when the sesqui-*TPS* gene was significantly down-regulated [30]. Likewise, a similar phenomenon appeared in our experiment, just because that besides having a basic sesquiterpene skeleton, the synthesis of dendrobine also requires a series of post-modification processes. To be specific, it could be speculated that the sharp increase in the content of dendrobine after 21 weeks may be the result of gene activation in the early stage, while the key enzyme genes in later stage were at a relatively low expression level, suggesting that the key enzyme genes of MVA pathway might be affected by dendrobine negative feedback regulation, to make sure that the content of dendrobine in plant could reach a relative balance.

Through the analysis of the results in the MEP pathway, we noticed that MEP pathway may not be directly involved in the regulation of dendrobine biosynthesis by MF23, but it had reported that these two ways are not completely separated [31]. For example, the addition of the *DXR* (a key enzyme gene in the MEP pathway) inhibitor not only inhibited the subsequent reaction of the MEP pathway, but also promoted the flow of IPP and DMAPP, the basic components of terpenoid synthesis, to the MVA pathway, which promoted an increase in the catalytic product of the MVA pathway without activating it [32–33]. Therefore, it may speculate that under the strong effect of MF23, the MEP pathway may get involved in the biosynthesis of dendrobine and cooperate with the MVA pathway to promote the biosynthesis of dendrobine.

### 3.5. Relationship between the key genes in downstream synthetic pathway and the content of dendrobine is clear on the bark medium

The downstream synthetic pathway of dendrobine mainly referred to the post-modification process of dendrobine. Researches had reported that the synthesis pathway of some secondary metabolite such as terpenoid, was inseparable from the effect of the modified enzymes[34]. For example, the synthesis pathway of paclitaxel included seven kinds of hydroxylation reactions and an epoxidation reaction [35]. So far, the key enzyme genes of the six hydroxylation reactions had been successfully cloned[36], and all of them showed good catalytic activity. In this study, through the exploration of the expression levels of the post-modification enzymes P450, methyltransferase and aminotransferase genes, we found that the genes were significantly up-regulated when dendrobine content increased. Nine weeks after inoculation, the corresponding relationship between the expression level of post-modification enzyme gene and the content of dendrobine is more significant. In particular, the aminotransferase *DAT* was significantly up-regulated in the experimental group at the 9^th^, 15^th^ and 21^st^ week compared to the control group, and reached the maximum at the 21^st^ week as well as the highest content of dendrobine appeared. In general, post-modification enzyme gene may play an important role in the process of MF23 promoting the increase of the dendrobine content when it grew in a natural environment.

## 4. Material and Methods

### 4.1. Source of materials

The tissue culture seedlings of *D. nobile* were obtained from Chishui Xintian humantang Pharmaceutical Limited Company (Guizhou, China).

### 4.2. Biological materials and culture conditions

The chestnut bark was washed, soaked for 24 hours, cut into pieces, and 70g of which was placed into each medium bottle, 80 ml deionized water was added, then autoclaved at 121°C for 180 minutes. Tissue culture seedlings of *D. nobile* with 3-5cm in height were cultured on bark medium, an artificial primary environment, which was similar to its primordial surroundings. Mycelial plugs from 20-day-old MF23 grew on the potato dextrose agar medium were placed in the root of the *D. nobile*, covered with bark. Then, the culture bottles were kept in a conventional greenhouse with 12 hours light and an illumination intensity of 2000LX, the temperature was controlled at 24 to 26°C, the plants without mycelial plugs were regarded as a control.

Plant samples were collected at different period: 1, 3, 6, 9, 12, 15, 18, 21, 24 weeks (**Figure S4**) the control group and the experimental group. 12 repetitions were carried on at each point of time, and the samples were selected randomly. All samples were divided into two portions; ones were used for morphological and chemical research, and the others were frozen in liquid nitrogen and stored for the RNA extraction.

### 4.3. Cultivation condition of D. nobile

One culture bottle with 12 plants as a repetition. We recorded the growth change of *D. nobile* such as the number of roots and stem diameter at each point of time. The fresh stems were weighed and parched at 55°C.

### 4.4. Determination of dendrobine content of D. nobile

Standard substances of dendrobine and the internal standard naphthalene were bought from Sinopharm Chemical Reagent Co, Ltd and Beijing Bei Na Chuang Lian Biotechnology Institute, respectively. The sample preparation for the GC analysis were performed according to the pharmacopoeia of China (2015). Chromatography was performed on an Agilent 6890 GC-FID with DB-1 capillary column (0.25 μm × 0.25 mm × 30 m) and nitrogen as a carrier gas.

Three replications were carried out to ensure the reliability of experimental results. 1μL of derivatized sample was injected, and components of the total ion chromatogram were extracted by the flame ionization detection. The linear regression equation y=0.1861x−0.1474 (r=0.9995) implied that the dendrobine concentration was linear, with a peak area in the range of 1.1~11 mg/L.

### 4.5. Light microscopy examination

Fixing fresh roots of *D. nobile* in formalin-acetic-acid-alcohol (FAA) followed as the approach described by Feder and O’Brien [37]. We dehydrated the samples in a graded ethanol series, then embedded in paraffin, stained with safranine and fast green, sealed with Gel Damar and then observed and take pictures on a light microscope matched with a camera (ZEISS Axio Imager A1).

### 4.6. RNA extraction

RNeasy Plant Mini Kit (cat. Nos 74903 and 74904) (Qiagen, Hilden, Germany) with the manufacturer’s instructions was used to extract the total RNA from samples of *D. nobile*. The conditions of Total RNA’s degradation and contamination were verified by electrophoresis in a 1.0% agarose gel in 0.5 × TBE (44.5 mM Tris–HCl, 44.5 mM boric acid and 1.25 mM Na2EDTA). The Qubit RNA Assay Kit in a Qubit 2.0 Fluorometer (Life Technologies, Carlsbad, CA, USA) was used to check the concentration of total RNA. In addition, the RNA Nano 6000 Assay Kit, equipped with a Bioanalyzer 2100 system (Agilent Technologies, Santa Clara, CA, USA), was used to assess the integrity of RNA.

### 4.7. Construction of cDNA library of D. nobile

The stem samples of the two groups of *D. nobile* at 9^th^ week and 21^st^ week respectively were picked out to extract RNA. An average of 1.5 μg RNA extracted from each sample, verified with good quality, was used as the input material for RNA sample preparations. Then, 8 samples with RNA integrity number (RIN) values above 8 were used for library construction, 2 repetitions were performed per treatment. The specific process of RNA extraction and other operation were introduced in the reference[20]that our research group had already published.

### 4.8. Confirmation of the infection-responsive expression profiles by qRT-PCR

The key enzyme genes involved in the formation of dendrobine including the mevalonate (MVA) pathway (**Table S1**), the methyl-D-erythritol 4-phosphate (MEP) pathway (**Table S2**) and the post-modification pathway (**Table S3**) were picked out from the transcriptome library created by our group.

The transcriptome library of *D. nobile* has already been established. Differential genes (DEGs) selected out were validated with qRT-PCR. The qRT-PCR was performed with the SYBR. Premix ExTaqTM (TaKaRa, Dalian, China) on an ABI 7500 Real-Time PCR System (Applied Biosystems, Foster City, CA, USA). The actin gene of *D. nobile* was used as a reference control [38]. The reaction was performed using the following conditions: denaturation at 95 °C for 30 s, followed by 40 cycles of amplification (95 °C for 5 s, 60 °C for 34 s). For each sample, three technical replicates of the qRT-PCR assay were used with a minimum of three biological replicates. Using the 2^−ΔΔCt^ method [39] to deal with the results and evaluate the expression of genes.

### 4.9. Statistical analysis of the data

Statistical software such as GraphPad Prism 5.0 and SPSS 19.0 were used to deal with the data of results. An unpaired t-test for the values was performed at P < 0.05

## 5. Conclusions

The results of content determination and microscopic observation showed that the fungus MF23 had a significant effect on the *D. nobile* grew on the bark medium. Its survival rate has been significantly increased in the later period during the inoculation. In addition, it found that the infection of MF23 was gradually progressed inwards, from root to exodermis and then to cortex through the microscopic observation. The expression levels of post-modification enzymes genes like cytochrome P450, aminotransferase and N-methyltransferase were studied, and it showed that these enzymes genes had a significant relevance with the content of dendrobine, especially the *DAT*, an aminotransferase enzyme gene, which was positively correlated with dendrobine biosynthesis. It implied that post-modification enzyme genes might play an important role in stimulating biosynthesis of dendrobine with the effect of MF23. The research laid an important foundation for MF23-induced increase in dendrobine content of *D. nobile*, and also provided a scientific basis for regulating the biosynthesis of dendrobine by molecular means. In conclusion, it reveals the proper mechanism of the mycorrhizal symbiosis mechanism in the wild, and lays a foundation for the possibility of using mycorrhizal fungi to promote the production of *D. nobile* in the field.

What is the most important is that the research we have carried out strongly suggested that MF23 was beneficial to the growth of *D. nobile* on the bark medium that close to the natural state, indicating that the endophytic fungus—MF23 is suitable for field production applications.

Simultaneously, further exploration such as the role and molecular mechanism of MF23 in improving the quality of *D. nobile* on the basis of experimental fields should be carried out in the future. Focusing on the basis of use of transgenic, RNAi and other technologies to verify the function of candidate genes. The study of protein will be performed to intuitively and accurately illuminate the molecular mechanism of how MF23 increase the effective component content of *D. nobile*. In short, the study carried out on the bark medium not only has important meaning in the application of the filed, but also lays the foundation on further researches.

## Declarations

### Ethics approval and consent to participate

Not applicable.

### Consent for publication

Not applicable.

### Availability of data and materials

The datasets used and/or analyzed during the current study are available from the corresponding author on reasonable request.

### Authors’ contributions

BL and SG conceived and designed the experiments. XX and QL performed the experiments, analyzed the data, and wrote the paper. GD provided technological guidance for the chemistry and edited the paper. All authors read and approved the final manuscript.

## Acknowledgements

We gratefully thank our reviewers for the suggestions and comments.

